# Genetic architecture of the white matter connectome of the human brain

**DOI:** 10.1101/2022.05.10.491289

**Authors:** Zhiqiang Sha, Dick Schijven, Simon E. Fisher, Clyde Francks

## Abstract

White matter tracts form the structural basis of large-scale functional networks in the human brain. We applied brain-wide tractography to diffusion images from 30,810 adult participants (UK Biobank), and found significant heritability for 90 regional connectivity measures and 851 tract-wise connectivity measures. Multivariate genome-wide association analyses identified 355 independently associated lead SNPs across the genome, of which 77% had not been previously associated with human brain metrics. Enrichment analyses implicated neurodevelopmental processes including neurogenesis, neural differentiation, neural migration, neural projection guidance, and axon development, as well as prenatal brain expression especially in stem cells, astrocytes, microglia and neurons. We used the multivariate association profiles of lead SNPs to identify 26 genomic loci implicated in structural connectivity between core regions of the left-hemisphere language network, and also identified 6 loci associated with hemispheric left-right asymmetry of structural connectivity. Polygenic scores for schizophrenia, bipolar disorder, autism spectrum disorder, attention-deficit hyperactivity disorder, left-handedness, Alzheimer’s disease, amyotrophic lateral sclerosis, and epilepsy showed significant multivariate associations with structural connectivity, each implicating distinct sets of brain regions with trait-relevant functional profiles. This large-scale mapping study revealed common genetic contributions to the structural connectome of the human brain in the general adult population, highlighting links with polygenic disposition to brain disorders and behavioural traits.

**One sentence summary:** Variability in white matter fiber tracts of the human brain is associated with hundreds of newly discovered genomic loci that especially implicate stem, neural and glial cells during prenatal development, and is also associated with polygenic dispositions to various brain disorders and behavioural traits.

## Introduction

Cognitive functions and behaviours are supported by dynamic interactions of neural signals within large-scale brain networks(*1*). Neural signals propagate along white matter tracts that link cortical, subcortical, and cerebellar regions to form the structural connectome(*2, 3*). White matter tracts also modulate neural signals and distribute trophic factors between connected regions(*4, 5*), helping to establish and maintain functional specialization of sub-networks. Various heritable psychiatric and neurological disorders can involve altered white matter structural connectivity, relating for example to cognitive deficits, clinical presentation or *recovery*(*6–10*). It is therefore of great interest to understand which DNA variants, genes and pathways affect white matter tracts in the human brain, as they are likely to influence cognitive and behavioural variability in the population, as well as predisposition to brain disorders.

Diffusion tensor imaging (DTI) enables *in vivo* non-invasive study of white matter in the brain(*11, 12*). This technique characterizes the diffusion of water molecules, which occurs preferentially in parallel to nerve fiber tracts due to constraints imposed by axonal membranes and myelin sheaths(*13, 14*). Metrics commonly derived from DTI, such as fractional anisotropy or mean diffusivity, reflect regional white matter microstructure and can index its integrity(*13–15*). In contrast, white matter tractography involves defining fiber tracts at the macro-anatomical scale, and computing connectivity strengths by counting the streamlines that link each pair of regions. Streamlines are constructed to pass through multiple adjacent voxels in DTI data, when the principal diffusion tensor per voxel aligns well with some of its direct neighbors(*16*). Tractography therefore produces subject-specific measures of regional inter-connectivity that are ideally suited for brain network-level analysis.

Recently, genome-wide association studies (GWAS) have reported that a substantial proportion of inter-individual variability in white matter microstructural measures can be explained by common genetic variants, with single nucleotide polymorphism (SNP)-based heritabilities ranging from 22% to 66%(*17, 18*). These studies also identified specific genomic loci associated with microstructural measures of white matter integrity(*17, 18*). However, to our knowledge, nerve fiber tractography has not previously been used for large-scale genome-wide association analysis of brain structural networks, likely due to heavy computational requirements for running tractography in tens of thousands of individuals.

Here, we aimed to characterize the genetic architecture of white matter structural network connectivity in the human brain, using fiber tractography. DTI data from 30,810 participants of the UK Biobank adult population dataset were used to construct the brain-wide structural connectivity network of each individual. In combination with genome-wide genotype data, we then carried out a set of genetic analyses of tractography-derived metrics, in terms of the sum of white matter connectivity linking to each of 90 brain regions (as network nodes), and each of 947 tract-wise measures that indicated connectivity between specific pairs of regions (as network edges). These analyses included SNP-based heritability estimation, multivariate GWAS (mvGWAS), and biological annotation of associated loci.

An important aspect of human brain organization is hemispheric specialization – the tendency of certain functions to be carried out dominantly by either the left or right cerebral hemispheres(*19*). Previous GWAS analyses have identified genetic loci associated with left-right asymmetries of cerebral cortical anatomy and/or handedness(*20–24*). Aspects of language function show especially strong lateralizations, with roughly 85% of people having left-hemisphere dominance(*25*). Such functional asymmetries may be partly underpinned by hemispheric asymmetries of white matter connectivity. Therefore, we used our brain-wide mvGWAS results to identify genomic loci that are associated with tract connectivities between core language-related regions of the left-hemisphere. In addition, for all structural connectivity metrics with paired left and right counterparts, we calculated their asymmetries and performed mvGWAS for any significantly heritable asymmetry measures, to identify genetic influences on hemispheric asymmetries of white matter connectivity.

Finally, we assessed how genetic disposition to brain disorders and other behavioural traits manifests in terms of white matter connectivity in the general population, and how this relates to cognitive processes. To do so, we mapped multivariate associations of white matter tractography metrics with polygenic scores for an array of heritable brain disorders and traits, including schizophrenia, bipolar disorder, autism, attention-deficit hyperactivity disorder, left-handedness, Alzheimer’s disease, amyotrophic lateral sclerosis, and epilepsy, and annotated the resulting brain maps with cognitive functions, using large-scale meta-analyzed functional neuroimaging data.

## Results

### White matter connectomes of 30,810 adults at regional and tract levels

For each of 30,810 adult participants with diffusion MRI and genetic data after quality control, we performed deterministic fiber tractography(*16*) between each pair of regions defined in the Automated Anatomical Labeling atlas(*26*) (45 regions per hemisphere comprising cerebral cortical and subcortical structures) (Fig. 1; Methods). In the structural connectivity matrix of each individual, each region was considered a node, and each tract considered an edge, with each tract comprising all streamlines that link a given pair of regions. We excluded tracts when more than 20% of individuals had no streamlines connecting a pair of regions, resulting in 947 tracts with streamline counts. To quantify the connectivity of each tract in each individual, streamline counts were divided by the individual-specific grey matter volume of the two regions that they connected, as larger regions tended to have more streamlines connecting to them. The volume-adjusted tract measures were also used to calculate the regional connectivity for each region (i.e. the sum of all edges connecting with a given node) within each participant. The resulting node and edge measures were adjusted for demographic and technical covariates, and normalized across individuals (Methods), before being used for the subsequent analyses of the study.

**Figure 1.**
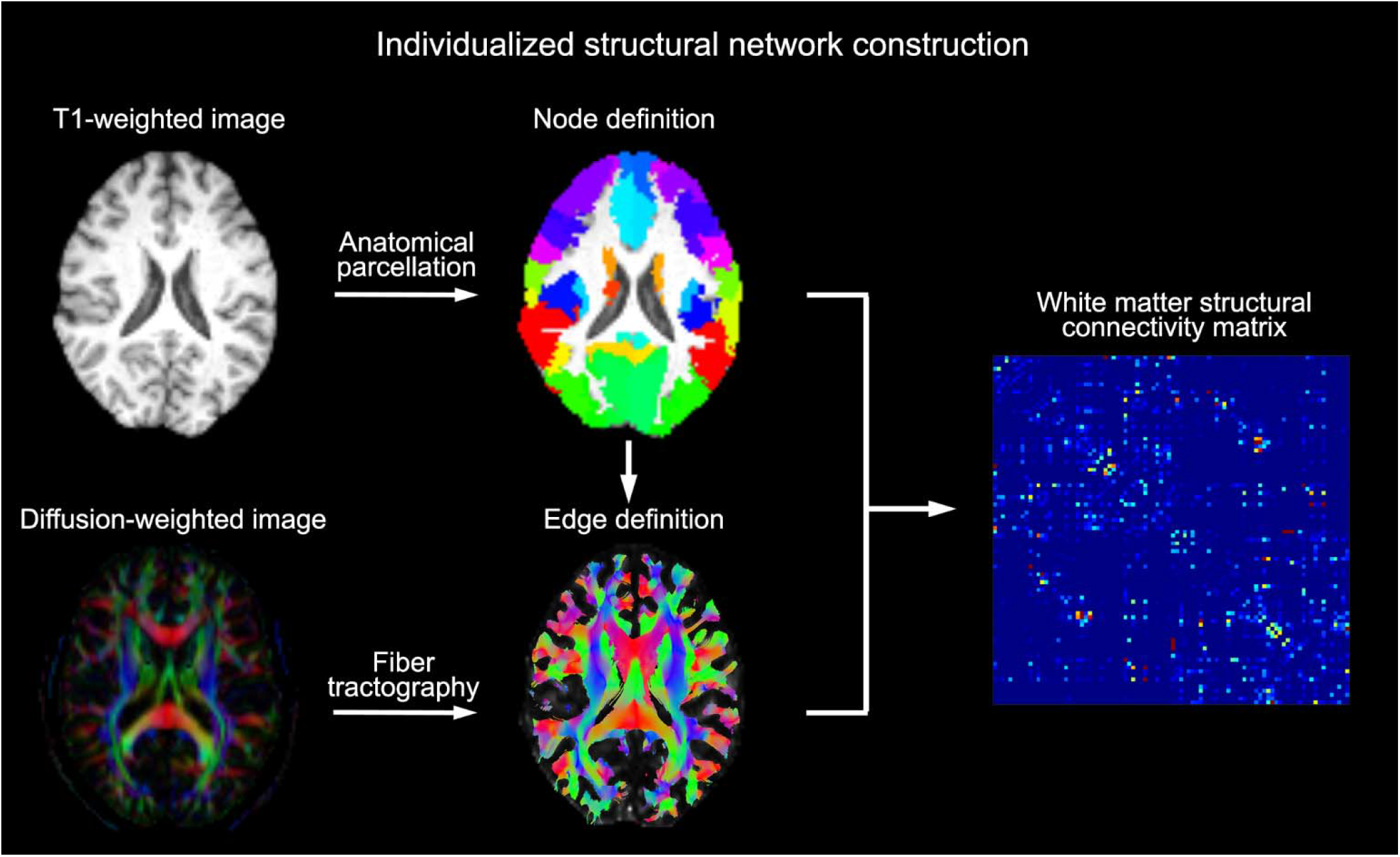
Schematic of white matter network construction within an individual brain. Network nodes were defined by mapping the Automated Anatomical Labeling atlas from common MNI space to individual space, with 45 regions per hemisphere (including cortical and subcortical structures). The edge between each pair of regions was defined as the number of streamlines constructed by tractography based on the corresponding diffusion tensor image, while adjusting for the volume of the connected regions. The process yielded a zero-diagonal symmetrical 90×90 undirected connectivity matrix for each of 30,810 participants (the upper triangles were then used for subsequent analyses).

Of the 947 tracts, 377 connected pairs of left-hemisphere regions, 355 connected pairs of right hemisphere regions, and 215 involved interhemispheric connections. The top 10% of regions in terms of connectivity included the supplementary motor cortex, precuneus, medial superior frontal cortex, and subcortical regions bilaterally – caudate and thalamus (Supplementary Figure 1 and Supplementary Table 1). The latter observation is consistent with previous studies showing that subcortical regions connect widely with the cerebral cortex, to generate reciprocal cortical-subcortical interactions that together support many cognitive functions(*4, 27, 28*).

### Heritabilities of region-wise and tract-wise connectivities

GCTA(*29*) was used to estimate the SNP-based heritability (*h*^2^) for each network measure, that is, the extent to which variance in each measure was explained by common genetic variants across the autosomes (Methods). For the 90 regional connectivities (i.e. connectivity of network nodes) all were significantly heritable (Bonferroni-corrected p<0.05), ranging from 7.8% to 29.5% (mean *h*^2^=18.5%; Fig. 2A and Supplementary Table 2). Most of the homologous regions in the left and right hemispheres showed similar heritabilities, while some regions showed prominent differences, such as the inferior parietal cortex (left: 27.0% vs. right: 19.42%), pars triangularis (left: 23.4% vs. right: 16.9%) and inferior occipital cortex (left: 8.0% vs. right: 15.7%; Fig. 2A and Supplementary Table 2). Eleven regional connectivities showed *h^2^* estimates >25% (Supplementary Table 1), with the superior temporal cortex in the left hemisphere being highest (*h*^2^=29.5%, p<1×10^−20^).

**Figure 2.**
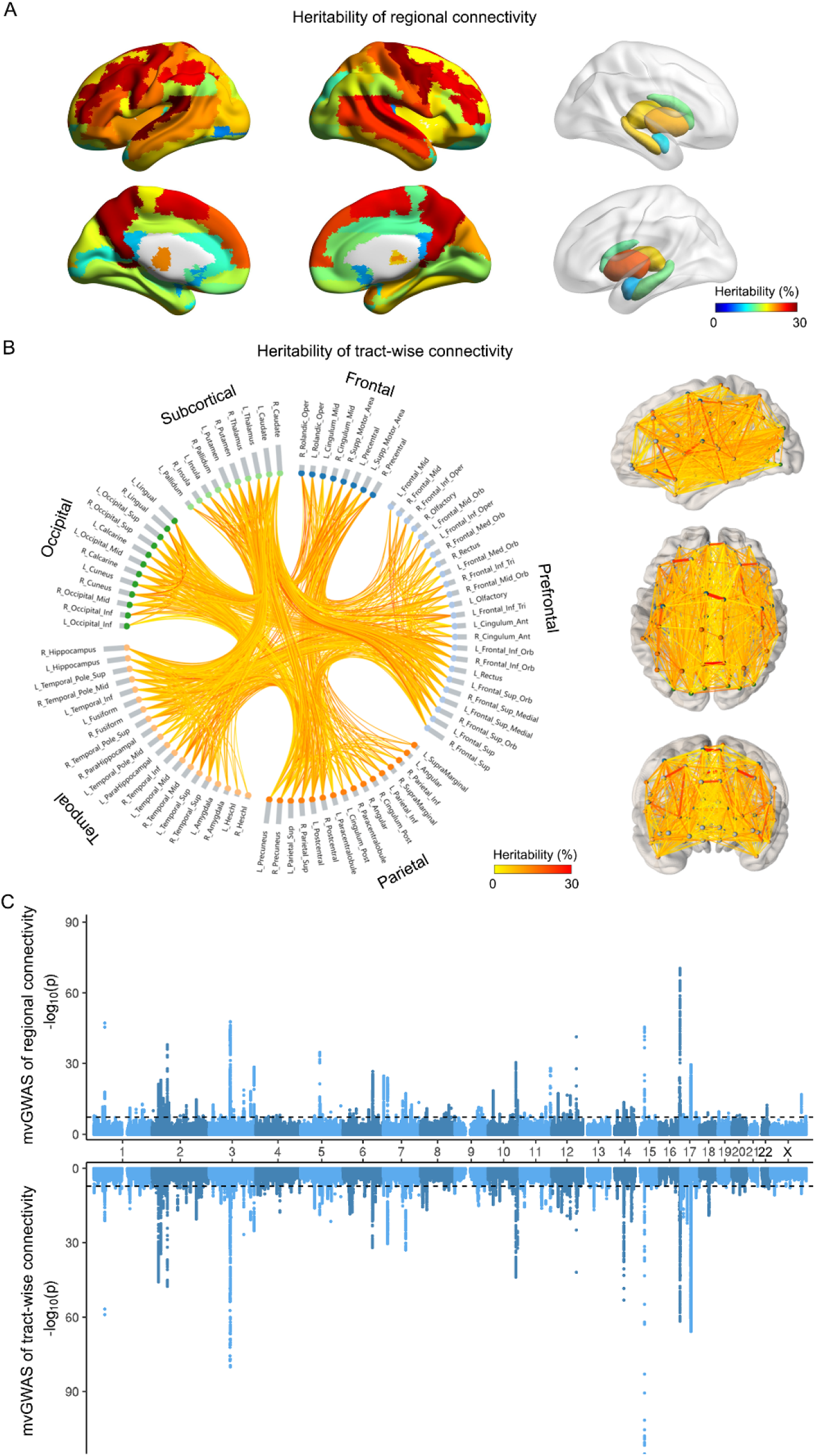
SNP-based heritability and multivariate GWAS analyses of regional connectivity and tract-wise connectivity in 30,810 participants. (A) All 90 regional connectivities showed significant SNP-based heritability after Bonferroni correction, ranging from 7.8% to 29.5%. (B) 851 out of 947 tract-wise connectivities showed significant SNP-based heritability after Bonferroni correction, ranging from 4.6% to 29.5%. Right panel: brain maps. Left panel: nodes grouped by frontal, prefrontal, parietal, temporal, and occipital cortical lobes, and subcortical structures. (C) Miami plot for multivariate GWAS of 90 regional connectivities (upper) and 851 tract-wise connectivities (lower). The black lines indicate the genome-wide significance threshold p<5×10^−8^ (Methods).

851 of 947 tract-wise connectivities (network edges) showed significant heritability (Bonferroni-corrected p<0.05), ranging from 4.6% to 29.5% (mean 9.6%), with a mean *h*^2^ of 9.9% for 351 tracts within the left hemisphere, a mean *h*^2^ of 10.0% for 333 tracts within the right hemisphere, and a mean *h*^2^ of 8.1% for 167 interhemispheric tracts (Fig. 2B and Supplementary Tables 3-5). Eleven out of 851 tract-wise connectivities had *h*^2^>20%, primarily for tracts linking bilateral frontal regions (e.g. superior and middle frontal cortex), supplementary motor and occipital cortex (e.g. cuneus and lingual).

We calculated the Euclidean distance between the centroids of each pair of connected regions (Methods), to index the relative physical distances between them. The heritabilities of tract-wise connectivities were negatively correlated with Euclidean distance across the 851 tracts (r=-0.14; Supplementary Figure 2), suggesting that shortrange tracts tend to be under stronger genetic control, and/or that they were more reliably measured.

### Multivariate genome-wide association analyses of regional and tract-wise connectivities

MOSTest(*30*) was used to perform two separate mvGWAS analyses, first for the 90 regional connectivity measures in a single multivariate genome-wide screen, and then for the 851 tract-wise connectivities in another single multivariate genome-wide screen, both times in relation to 9,803,735 SNPs spanning the genome. This analysis examined each SNP separately for its associations with multiple structural network measures, by simultaneously modelling the distributed nature of genetic influences across the brain (Methods). FUMA(*31*) was used to clump mvGWAS results on the basis of linkage disequilibrium (LD) and to identify independent lead SNPs at each associated genomic locus (Methods). At the 5×10^−8^ significance level, we identified 154 lead SNPs in 128 distinct genomic loci associated with regional connectivities (node level) (Fig. 2C, Supplementary Figure 3 and Supplementary Table 6), and 231 lead SNPs in 181 distinct genomic loci associated with tract-wise connectivities (edge level) (Fig. 2C, Supplementary Figure 3 and Supplementary Table 7). 97 genomic loci were found in common between the regional connectivity mvGWAS and tract-wise connectivity mvGWAS. Permutation analysis under the null hypothesis of no association indicated that MOSTest correctly controlled type I error (Methods; Supplementary Figures 4 and 5). Except for chromosome 21, each chromosome had at least one locus associated with either regional connectivity or tract-wise connectivity.

For each lead SNP, MOSTest indicated the contribution of each brain metric to its multivariate association, by reporting a z-score derived from each metric’s univariate association with that SNP (Methods; Supplementary Tables 8 and 9). In the regional connectivity (node level) mvGWAS, regions with the greatest magnitude z-scores considered across all lead SNPs were the bilateral middle frontal cortex (left mean |z|=2.02, right mean |z|=1.89), bilateral putamen (left mean |z|=2.01, right mean |z|=1.82), right superior frontal cortex (mean |z|=1.80) and middle cingulate cortex (mean |z|=1.78; Supplementary Figure 6 and Supplementary Table 10). For example, the left middle frontal cortex, which had the highest overall contribution across lead SNPs (mean |z|=2.02), was especially strongly associated with rs756705025 on 5q14.2 (z=8.90), rs67827860 on 5q14.3 (z=8.37), rs2696626 on 17q21.31 (z=7.11), rs55938136 on 17q21.31 (z=6.66) and rs4385132 on 4q12 (z=6.59; Supplementary Table 10).

In the mvGWAS of tract-wise connectivity (edge level), tracts that showed high magnitude z-scores considered across all lead SNPs mainly connected the precuneus, calcarine, middle temporal and pre- and post-central cortex (Supplementary Table 11 and Supplementary Figure 7). The tract linking the left and right precuneus had the greatest contribution across lead SNPs (mean |z|=1.59), and was especially associated with the variants rs946711 on 10p12.31 (z=-5.58) and 3:190646282_TA_T on 3q28 (z=-5.53).

### The majority of genomic loci associated with structural connectivity are novel

Together, our regional connectivity mvGWAS and tract-wise connectivity mvGWAS identified 355 lead SNPs, of which 105 were previously associated with at least one trait in the NHGRI-EBI GWAS catalog(*32*) (Supplementary Tables 6 and 7). This indicates that the majority (70.4%) of loci implicated here in the structural connectome were not identified by previous studies. There were 68 SNPs in common with those reported in previous GWAS of brain measures(*18, 30, 33–35*). Specifically, 48 of our lead SNPs were previously associated with brain regional volumes(*33, 36*), 30 with regional cortical thicknesses(*30, 35*), 35 with regional cortical surface areas(*35, 37*), and 20 with white matter microstructure(*18, 38*). Apart from brain measures, 11 of our lead SNPs were associated with mental health traits (e.g. autism, schizophrenia, and anxiety)(*39, 40*), 12 of our lead SNPs with cognitive functions (e.g. cognitive ability and performance)(*41, 42*), 4 of our lead SNPs with neurological diseases (e.g. Alzheimer’s disease and epilepsy)(*43, 44*), and 42 of our lead SNPs with non-brain physiological and physical variables (e.g. waist-hip ratio, cholesterol levels and lung function)(*45, 46*). In addition, we compared our results with those reported in a recent GWAS of white matter microstructure integrity for which the results have not been deposited in the GWAS Catalog(*17*): 33 of their lead SNPs overlapped with those from our mvGWAS analyses (Supplementary Table 12).

### Functional annotations of genomic loci associated with the structural connectome

We used FUMA(*31*) to annotate SNPs to genes at significantly associated loci by three strategies: physical position, expression quantitative trait locus (eQTL) information and chromatin interactions (Methods). For the regional connectivity mvGWAS (node level), 960 unique genes were identified through these three strategies (Supplementary Table 13 and Supplementary Figure 8). 101 out of 154 lead SNPs had at least one eQTL or chromatin interaction annotation, indicating that these variants (or other variants in high LD with them) affect gene expression. For example, rs7935166 on 11p11.2 (multivariate z=5.71, p=1.15×10^−8^) is intronic to *CD82*, which has been reported to promote oligodendrocyte differentiation and myelination of white matter(*47*). This lead SNP is not only a brain eQTL(*48, 49*) of *CD82*, but also shows evidence for cross-locus chromatin interaction via the promoter of *CD82* in adult brain(*48*). Another example: rs35396874 on 6q21 (multivariate z=6.64, p=3.17×10^−11^) affects the expression of its surrounding gene *FOXO3*, a core element of the TLR/AKT/FoxO3 pathway that is important for repairing white matter injury mediated by oligodendrocyte progenitor cell differentiation(*50, 51*).

For the tract-wise connectivity mvGWAS (edge level), functional annotation identified 1530 unique genes (Supplementary Table 14 and Supplementary Figure 8). 148 of 231 lead SNPs had at least one eQTL annotation or chromatin interaction. For example, rs13084442 on 3q26.31 (multivariate z=6.34, p=2.32× 10^−10^) is an eQTL(*52*) of *TNIK*, a gene associated with neurogenesis and intellectual disability(*53, 54*). The same lead variant is also located in a region having a chromatin interaction with *TNIK* promoters in fetal and adult cortex(*55*). Similarly, the SNP rs28413051 on 4q31.23 (multivariate z=6.28, p=3.47×10^−10^) is an eQTL of *DCLK2* that is important for axon growth cone formation and neural migration(*56, 57*), and is also within a region interacting with the promoter of *DCLK2* in neural progenitor cells(*50*). A further example: allele C of rs13107325 on 4q24 (multivariate z=5.77, p= 7.99×10^−9^ in the regional connectivity mvGWAS and multivariate z=8.74, p=2.37×10^−18^ in the tract-wise connectivity mvGWAS) is a missense coding variant in the gene *SLC39A8* that showed a high combined annotation-dependent depletion (CADD) score of 23.1 (Methods), which indicates that this SNP is deleterious (its frequency was 7.01%). The same SNP has been associated with white matter microstructure integrity(*34*), schizophrenia(*58, 59*) and children’s behavioural problems(*60*).

### Gene-based association analysis and gene set enrichment analysis for the brain’s structural connectome

We used MAGMA(*37*) to perform gene-based association analysis, which combines the mvGWAS evidence for association at each SNP within a given gene, while controlling for LD.

For regional connectivities we identified 343 significant genes after Bonferroni correction (Supplementary Table 15 and Supplementary Figure 9), 255 of which overlapped with those annotated by at least one of the three strategies used above (i.e. physical location, eQTL annotation or chromatin interaction). The gene-based p values were then used as input to perform gene-set enrichment analysis, in relation to 15,488 previously defined functional sets within the MSigDB database(*61*). Sixty-one gene sets showed significant enrichment (Bonferroni adjusted p<0.05, Fig 3A and Supplementary Table 16) which mainly implicated neurodevelopmental processes, such as “go_neurogenesis” (beta=0.18, p=5.53×10^−13^; the most significant set), “go_neuron_differentiation” (beta=0.18, p=1.55×10^−10^), and “go_cell_morphogenesis_involved_in_neuron_differentiation” (beta=0.25, p=3.39×10^−10^).

**Figure 3.**
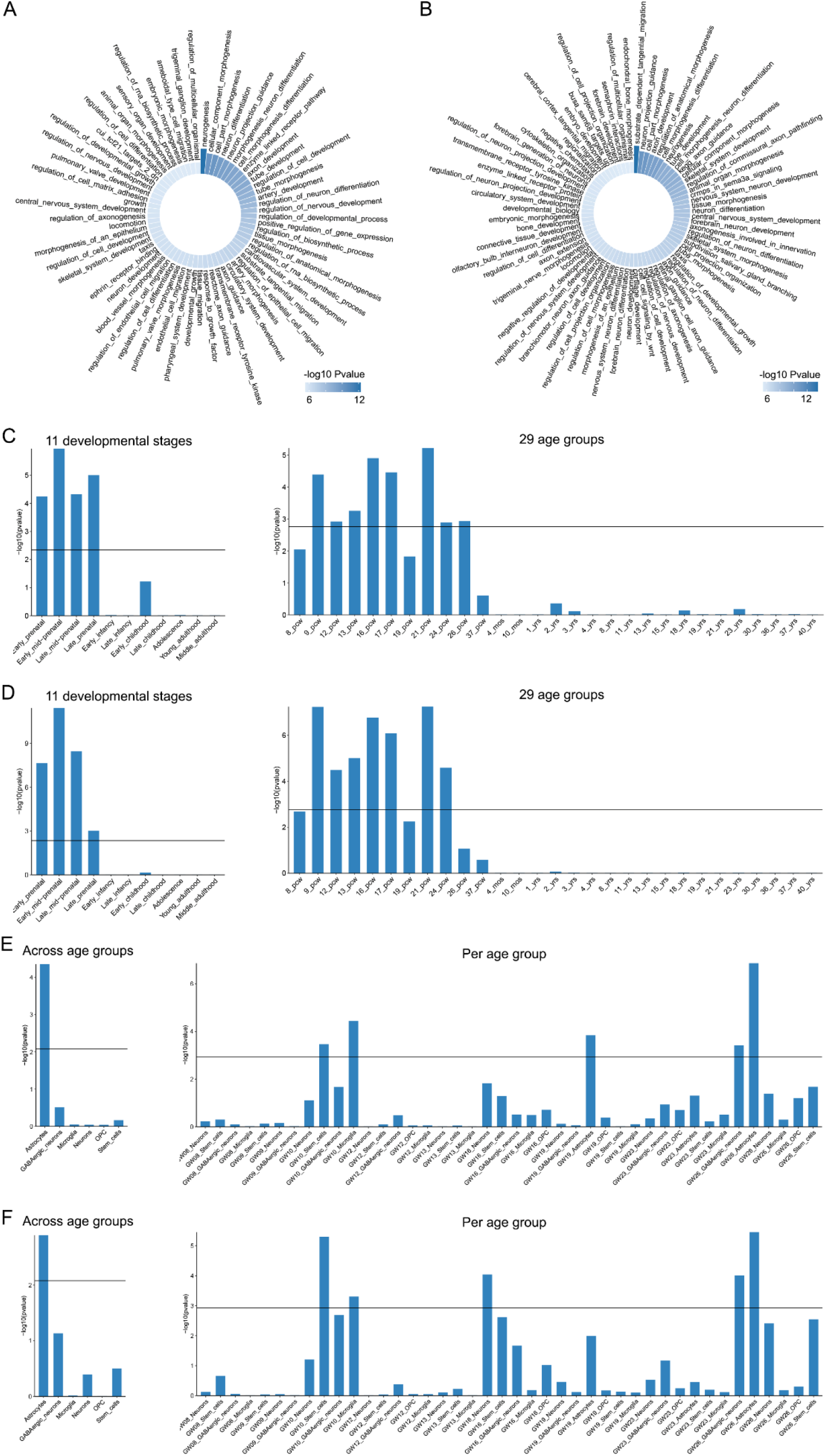
Genes associated with variation in the adult white matter connectome are enriched for specific neurodevelopmental roles. (A) Sixty-one functionally-defined gene sets showed significant enrichment of association with regional connectivity. (B) Seventy-two functionally-defined gene sets showed significant enrichment of association with tract-wise connectivity. (C-D) Based on BrainSpan data from 11 lifespan stages or 29 age groups, genes associated with variation in (C) adult regional connectivity and (D) adult tractwise connectivity show upregulation in the human brain prenatally. (E-F) Based on single-cell gene expression data from the prenatal brain, genes associated with variation in (E) adult regional connectivity show upregulation in astrocytes when considering all prenatal age groups combined, and in stem cells and microglia at 10 gestational weeks, astrocytes at 19 gestational weeks, and GABAergic neurons and astrocytes at 26 gestational weeks when breaking down by developmental stages, and similarly genes associated with variation in (F) adult tract-wise connectivity show upregulation in astrocytes when considering all prenatal age groups combined, and in stem cells and microglia at 10 gestational weeks, neurons at 16 gestational weeks, and GABAergic neurons and astrocytes at 26 gestational weeks when breaking down by developmental stages. (C-F) Black lines indicate the significance threshold p<0.05 after Bonferroni correction. PCW: post-conceptional weeks.

For tract-wise connectivities we identified 618 genes with significant gene-based association (Bonferroni-corrected p values <0.05), 461 of which overlapped with genes mapped through physical location, eQTL annotation or chromatin interaction (Supplementary Table 17 and Supplementary Figure 9). Seventy-two gene sets were significant after Bonferroni correction (Fig. 3B and Supplementary Table 18) related especially to neural migration and the development of neural projections, such as “go_substrate_dependent_cerebral_cortex_tangential_migration” (beta=3.98, p=2.61×10^−14^; the most significant set), “go_neuron_projection_guidance” (beta=0.41, p=8.59×10^−12^), and “go_axon_development” (beta=0.29, p=3.45×10^−11^).

We tested our genome-wide, gene-based p values with respect to human brain gene expression data from the BrainSpan database(*62*), grouped according to 11 lifespan stages or 29 different age groups. Genes associated with regional connectivity showed upregulation on average across much of the prenatal period, ranging from early (beta=0.04, p=5.84×10^−5^) to late (beta=0.08, p=1.01×10^−5^) prenatal stages, or from 9 (beta=0.002, p=4.15×10^−5^) to 26 (beta=0.003, p=1.18×10^−3^) post-conceptional weeks (Bonferroni-corrected p values <0.05; Fig. 3C and Supplementary Table 19). Similarly, genes associated with tract-wise connectivities showed upregulation on average during early (beta=0.06, p=2.35×10^−8^) to late (beta=0.06, p=1.01×10^−3^) prenatal stages, or from 9 (beta=0.003, p=5.92×10^−8^) to 24 (beta=0.003, p=2.67×10^−5^) post-conceptional weeks (Bonferroni-corrected p values <0.05; Fig. 3D and Supplementary Table 20).

We also examined our genome-wide, gene-based association p values with respect to two independent single-cell gene expression datasets derived from human prefrontal cortex samples of different ages (GSE104276)(*63*). Combining across age groups, average upregulation was observed in astrocytes for genes associated with both regional connectivity (beta=0.05, p=4.34×10^−5^) and tract-wise connectivity (beta=0.04, p=1.27×10^−3^) (Bonferroni-corrected p values <0.05; Fig. 3E and 3F and Supplementary Tables 21 and 22). Breaking down by age, genes associated with regional connectivity were upregulated on average in microglia (beta=0.02, p=3.72×10^−5^) and stem cells (beta=0.05, p=3.51×10^−4^) at 10 gestational weeks of age (GW), astrocytes at 19GW (beta=0.02, p=1.48×10^−4^) and 26GW (beta=0.05, p=1.35×10^−7^), and GABAergic neurons at 26GW (beta=0.04, p=3.97×10^−4^) (Fig. 3E and Supplementary Table 21). Similarly, genes associated with tract-wise connectivities (edge-wise) showed upregulation on average in microglia (beta=0.02, p=5.07×10^−4^) and stem cells (beta=0.06, p=5.18×10^−6^) at 10GW, neurons at 16GW (beta=0.06, p=9.32×10^−5^), and astrocytes (beta=0.05, p=3.50×10^−6^) and GABAergic neurons at 26GW (beta=0.05, p=1.01×10^−4^); Fig. 3F and Supplementary Table 22).

### Genetics of left-hemisphere language network connectivity

We selected four left-hemisphere regions that correspond to a network that is reliably activated by sentence-level language tasks in a left-lateralized manner in the majority of people and across languages(*64–67*), i.e. the opercular and triangular parts of inferior frontal cortex (Broca’s region), and the superior and middle temporal cortex (including Wernicke’s region, Fig. 4). These four regions are linked by six tracts with heritabilities ranging from 7.3% to 17.1% (Supplementary Table 3). Of the 231 lead SNPs from our brain-wide mvGWAS of tract-wise connectivity, 26 were significantly associated with at least one of these six tracts according to the tract-specific z-scores derived from MOSTest (Bonferroni correction at 0.05; Supplementary Table 23). For example, rs12636275 on 3p11.1 is located within an intron of *EPHA3*, a gene that encodes an ephrin receptor subunit that regulates the formation of axon projection maps(*68*), and has also been associated with functional connectivity between language-related regions(*69*). Another example: rs7580864 on 2q33.1 is an eQTL of *PLCL1* that is implicated in autism(*70, 71*), a neurodevelopmental disorder that often affects language and social skills. Other positional candidate genes based on the 26 SNPs include *CRHR1*, encoding corticotropin releasing hormone receptor 1, and *CENPW* (centromere protein W) involved in chromosome maintenance and the cell cycle (Fig. 4 and Supplementary Table 23).

**Figure 4.**
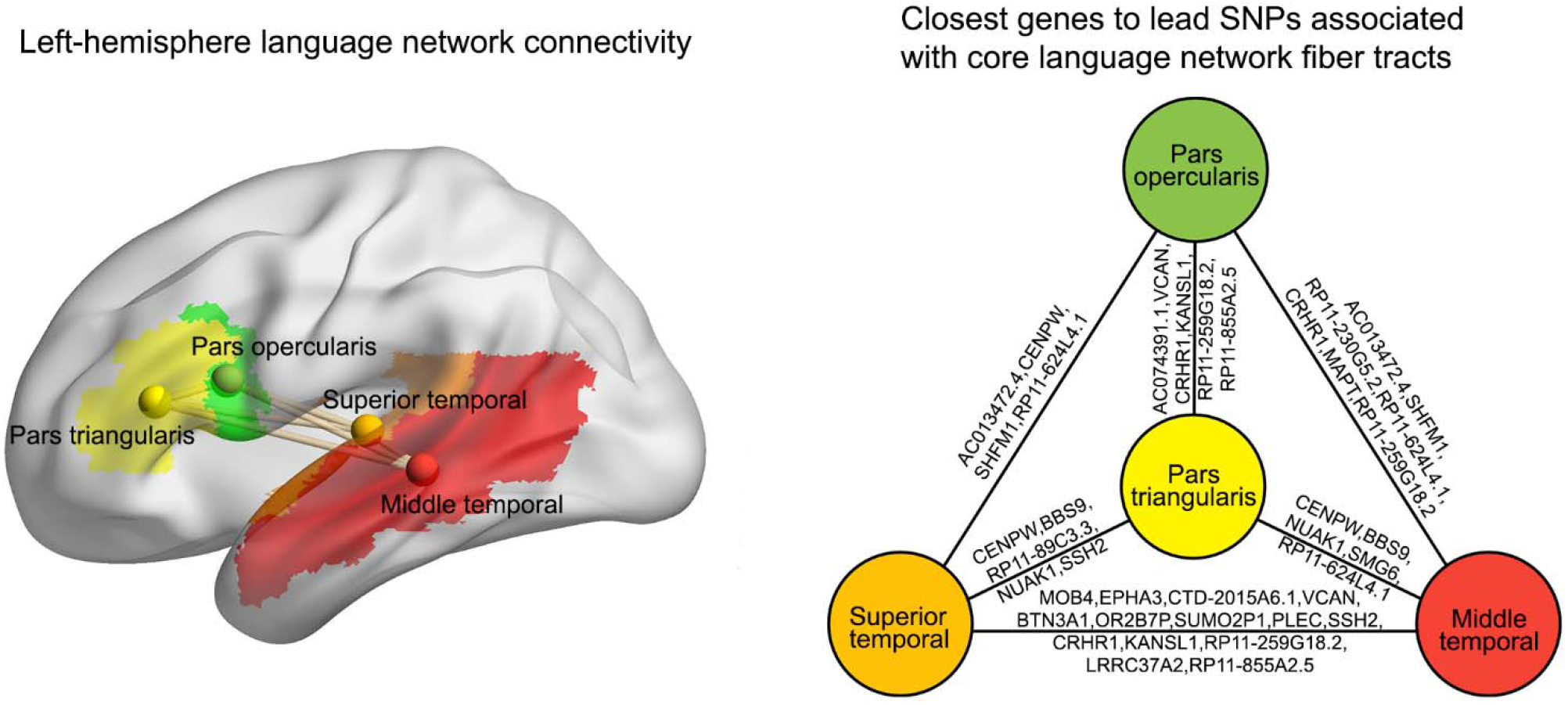
Genetics of left-hemisphere language network connectivity. Left: Four regions with core functions in the left-hemisphere language network, encompassing the classically defined Broca’s (frontal lobe) and Wernicke’s (temporal lobe) areas. Also shown are the six edges connecting these four regions when considered as network nodes. Right: The closest genes to independent lead SNPs from the brain-wide multivariate GWAS of tract-wise connectivity, that showed significant association with at least one of the six left-hemisphere language network tracts (Bonferroni correction at 0.05; Supplementary Table 23).

### Genetics of left-right asymmetry of the structural connectome

To investigate hemispheric specialization of structural connectivity in the brain, we computed the asymmetry index, AI=2(left-right)/(left+right), for each bilateral pair of regional connectivity and fiber tract metrics in each individual (Methods). For regional connectivity, all but one region showed population-level average asymmetry (mean AI different from 0, Bonferroni-corrected p<0.05), with regions comprising Broca’s area, plus the superior and middle temporal, postcentral, orbitofrontal and calcarine cortex, as well as the thalamus, showing strong leftward asymmetry of structural connectivity (Supplementary Figure 10 and Supplementary Table 24). Fifteen regional connectivity asymmetries were significantly heritable, mean *h*^2^=4.54% (range 3.82% to 5.34%, Bonferroni-corrected p<0.05; Supplementary Figure 11 and Supplementary Table 25), including asymmetries of structural connectivity linking to the dorsolateral prefrontal cortex, Broca’s area, supramarginal and fusiform cortex (Supplementary Figure 11).

At the level of tract-wise connectivity asymmetries, none showed significant population-level average asymmetry after multiple testing correction, but there were 107 tract-wise connectivity asymmetries with significant heritability, mean *h*^2^=6.75% (range 4.34% to 12.17%, Bonferroni-corrected p<0.05; Supplementary Figure 11 and Supplementary Tables 26-29).

We performed two further mvGWAS analyses, one for heritable regional connectivity asymmetries and one for heritable tract-wise connectivity asymmetries, using MOSTest(*30*). There were two independent lead SNPs in two distinct genomic loci associated with regional connectivity asymmetries at the 5×10^−8^ significance level (Supplementary Figures 11 and 12 and Supplementary Table 30): rs28520337 on 15q14 (multivariate z=7.11,p=1.14×10^−12^) is located in an intron of *RP11-624L4.1* and its multivariate association was driven particularly by the asymmetries of rolandic operculum (z-score=5.06) and supramarginal gyrus (z-score=6.47) connectivity (Supplementary Table 31). This SNP is in high LD with rs4924345 (r^2^=0.76), previously associated with cortical surface area(*35*). The second lead SNP rs56023709 on 16q24.2 (multivariate z=6.15, p=7.92×10^−10^) is located in *C16orf95* and its multivariate association was driven particularly by the asymmetries of supplementary motor cortex (z-score=-4.58) and pallidum (z-score=-3.63) connectivity. This SNP is in high LD with rs12711472 (r^2^=0.99), previously associated with thalamus volume(*30*).

For tract-wise connectivity asymmetries, mvGWAS identified four independent lead SNPs in four distinct genomic loci at the 5×10^−8^ significance level (Supplementary Figures 11 and 13, Supplementary Table 32): rs73219794 on 4p15.1 (multivariate z=5.66, p=1.54×10^−8^; nearest gene *RP11-180C1.1*), rs182149107 on Xp21.2 (multivariate z=5.77, p=7.87×10^−9^; nearest gene *IL1RAPL1*, encoding a synaptic adhesion molecule, mutated in intellectual disability and autism(*72*)), rs4824483 on Xp11.23 (multivariate z=5.80, p=6.56×10^−9^; downstream of *GAGE1*, upregulated in some glioblastomas(*73*)) and rs140894649 on Xq11.2 (multivariate z=6.61, p=3.72×10^−11^; nearest gene *MTMR8*, a phosphatase enzyme that regulates actin filament modeling(*74*) – consistent with possible roles of cytoskeleton-related genes in patterning left-right asymmetry(*20, 21, 75*)). None of these SNPs had association z-scores that stood out for any particular tract connectivity asymmetries (Supplementary Table 33), i.e. contributions to their significant multivariate associations were widely distributed across many tract asymmetries.

Gene-based association analyses for regional connectivity asymmetries or tract-wise connectivity asymmetries did not identify significantly associated genes, and neither did these gene-based association statistics show significant enrichment with respect to functionally defined gene sets, or differential expression levels in brain tissue at specific lifespan stages or cell types in the prenatal brain (Methods; Supplementary Tables 34-37).

### Multivariate associations of the structural connectome with polygenic scores for brain disorders and behavioural traits

For each of the 30,810 individuals in our study sample we calculated polygenic scores(*76*) for various brain disorders or behavioural traits that have shown associations with white matter variation, using previously published GWAS summary statistics: schizophrenia(*17, 77–79*), bipolar disorder(*80–82*), autism(*17, 20, 78, 83*), attention deficit/hyperactivity disorder(*84–86*), left-handedness(*21, 23*), Alzheimer’s disease(*87–89*), amyotrophic lateral sclerosis(*90–92*), and epilepsy(*93–95*) (Methods). There were 18 significant partial correlations (i.e. adjusted for confounds including sex and age – see Methods) between different pairs of these polygenic scores across individuals (Bonferroni-corrected p<0.05): 16 correlations were positive, with the highest between polygenic scores for schizophrenia and bipolar disorder (r=0.36, p<1×10^−200^), and between attention deficit/hyperactivity disorder and autism (r=0.33, p<1×10^−200^), while 2 were negative, between polygenic scores for amyotrophic lateral sclerosis and bipolar disorder (r=-0.03, p=2.26× 10^−6^), and amyotrophic lateral sclerosis and autism (r=-0.03, p=8.81×10^−6^; Supplementary Table 38 and Supplementary Figure 14).

Separately for each of these polygenic scores, we used canonical correlation analysis to investigate their multivariate associations with the 90 heritable regional connectivity measures across the 30,810 individuals. All canonical correlations were highly significant: schizophrenia r=0.07, p=8.98× 10^−34^; bipolar disorder r=0.07, p=1.53×10^−35^; autism r=0.06, p=7.87×10^−24^; attention deficit/hyperactivity disorder r=0.08, p=7.84×10^−44^; left-handedness r=0.07, p=1.74×10^−31^; Alzheimer’s disease r=0.07, p=4.14×10^−33^; amyotrophic lateral sclerosis r=0.06, p=1.29×10^−25^; epilepsy r=0.05, p=1.49× 10^−20^. Therefore, polygenic dispositions to these various disorders or behavioural traits in the general population are partly reflected in the brain’s white matter connectivity.

Canonical correlation analyses yielded loadings for each regional connectivity measure, reflecting the extent and direction of each measure’s association with polygenic disposition for a given disorder/behavioural trait. For psychiatric disorders the majority of loadings were negative, i.e. increased polygenic risk for these disorders was more often associated with reduced than increased connectivity across regions (Fig. 5, Supplementary Table 39). This was especially marked for polygenic risks for schizophrenia (85 regions with negative loadings, 5 regions with positive loadings), bipolar disorder (81 negative, 9 positive) and autism (64 negative, 26 positive). Polygenic disposition to left-handedness was also associated with more reduced regional connectivities (62 negative loadings) than increased regional connectivities (28 positive loadings). In contrast, increased polygenic risk for Alzheimer’s disease was associated with increased white matter connectivity for a majority of brain regions (62 out of 90) in the UK Biobank data, even while some core regions of disorder pathology showed decreased connectivity, such as posterior cingulate and medial temporal cortex(*96–98*). (These results remained stable when excluding the *APOE* locus that is known to have a substantial individual effect on Alzheimer’s disease risk – see Methods and Supplementary Table 40.) Similar observations were made for polygenic risk for amyotrophic lateral sclerosis, where 74 out of 90 regions showed positive loadings (Fig. 5, Supplementary Table 39).

**Figure 5.**
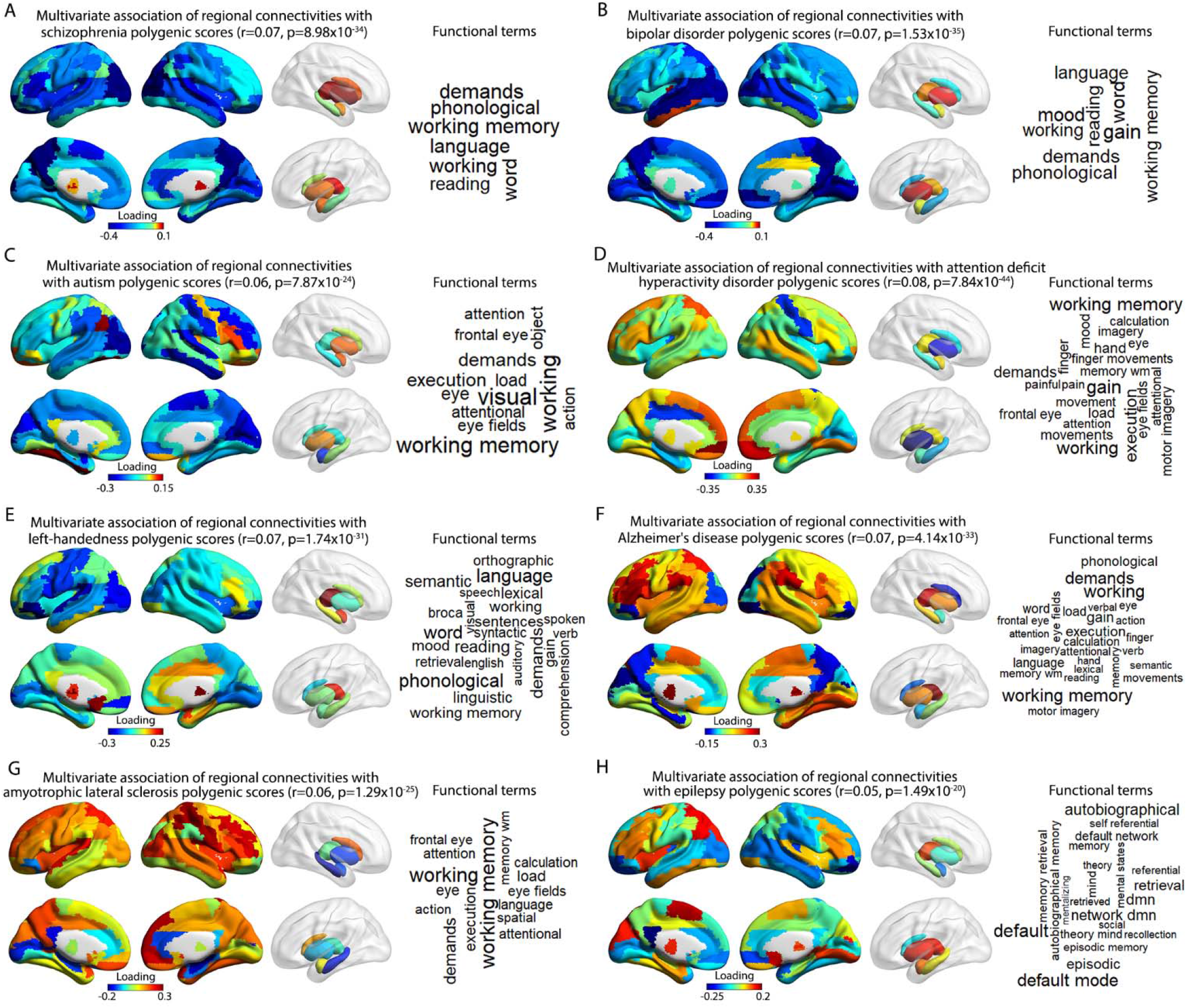
Polygenic dispositions to various brain-related disorders or behavioural traits show multivariate associations with regional white matter connectivities in 30,810 participants. Loadings are shown from canonical correlation analyses that indicate the extent and direction to which each regional connectivity is associated with polygenic scores for (A) schizophrenia, (B) bipolar disorder, (C) autism, (D) attention deficit/hyperactivity disorder, (E) left-handedness, (F) Alzheimer’s disease, (G) amyotrophic lateral sclerosis and (H) epilepsy. A positive loading (red) indicates a higher regional connectivity associated with increased polygenic disposition for a given disorder/behavioural trait, while a negative loading (blue) represents a lower regional connectivity associated with increased polygenic disposition for a given disorder/behavioural trait. Word clouds represent functions associated with the map of regions showing the strongest loadings (|r|>0.2) for each polygenic score. Functions were assigned using large-scale meta-analyzed functional neuroimaging data (Methods). The font sizes in the word clouds represent correlation magnitudes between the meta-analyzed functional maps for those terms and the co-activation map for the set of regions associated with each polygenic score. See Supplementary Table 41 for the correlation values.

For each polygenic score we identified the specific regional connectivities that showed the strongest loadings in canonical correlation analyses, i.e. regions with loadings >0.2 or <-0.2. These regions were used to create a single brain mask for each polygenic score, which was then used to query the Neurosynth database of 14,371 functional brain imaging studies(*99*). In this process, a brain-wide co-activation map was generated for each mask, based on all functional maps in the database, and these were then correlated with cognitive and behavioural term-specific maps derived from the studies included in the database(*99*).

For example, the mask for schizophrenia polygenic risk comprised 32 regions showing the strongest associations with white matter connectivity, distributed in the bilateral temporal, dorsoventral and posterior cingulate cortex (Fig. 5A and Supplementary Table 39), and there were 7 functional term-based correlations >0.2 with the corresponding co-activation map for these regions (Fig. 5A, Supplementary Figure 15 and Supplementary Table 41), including ‘working memory’ and ‘language’. This suggests that polygenic disposition to schizophrenia influences the connectivity of brain regions especially involved in working memory and language (see Discussion). The mask for bipolar disorder polygenic risk comprised 30 regions, including temporal, medial frontal, superior parietal, and visual cortex, as well as hippocampus and caudate, and these regions together received functional annotations of ‘mood’, ‘working memory’ and language-related processes (Fig. 5B, Supplementary Figure 15 and Supplementary Tables 39 and 41). Polygenic risk for autism was mainly associated with white matter connectivity of the right dorsolateral prefrontal, right temporal, right sensorimotor and bilateral visual cortex, as well as the left amygdala, and these regions were annotated with visual, working memory, executive and attention functions (Fig. 5C, Supplementary Figure 15 and Supplementary Tables 39 and 41). Polygenic disposition to left-handedness was associated with regional connectivity of Broca’s area, left superior temporal cortex, left medial prefrontal and left visual cortex and right thalamus, functionally annotated with language-related cognitive functions (Fig. 5E, Supplementary Figure 15 and Supplementary Tables 39 and 41). See Fig. 5, Supplementary Figure 15 and Supplementary Tables 39 and 41 for the equivalent maps and functional annotations for all disorder/trait polygenic scores.

The polygenic risks for bipolar disorder and schizophrenia had the most similar brain maps, in terms of regional structural connectivity associated with each of these polygenic risks (r=0.56 between the loadings for these two polygenic scores, across the 90 regions; Supplementary Figure 16 and Supplementary Table 42).

## Discussion

This large-scale mapping study employed white matter tractography and multivariate analysis to characterize the contributions of common genetic variants to individual differences in structural connectivity of the adult human brain. Multivariate associations between structural connectivity and polygenic dispositions to brain-related disorders or behavioural traits were also characterized, and described in terms of functional activations of the implicated brain regions. Together, these various analyses in over 30,000 individuals from the general population linked multiple levels of biological organization: from genes and cell types through developmental stages to adult brain structure and function, behaviour and individual differences, newly implicating hundreds of genomic loci.

Different brain regions are inter-connected through white matter nerve fibers - this fundamental property sub-serves functional networks involved in cognition and behaviour. In over 30,000 adults from the general population we found that inter-individual variation in white matter connectivity is especially influenced by genes that are i) active in the prenatal developing brain; ii) upregulated in stem cells, astrocytes, microglia, and neurons of the embryonic and fetal brain; and iii) involved in neurodevelopmental processes including neural migration, neural projection guidance and axon development. A likely neurodevelopmental origin of much inter-individual variation of adult white matter connectivity is consistent with findings from large-scale imaging genetics studies of other aspects of brain structural and functional variation(*17, 20, 35*). These statistical enrichment findings serve as a strong biological validation of the multivariate GWAS findings, as there was no reason for such clearly relevant functional enrichment to occur by chance in relation to brain white matter tracts.

Astrocytes are the largest class of brain glial cells with a range of known functions, including neuronal homeostasis and survival, regulation of synaptogenesis and synaptic transmission(*100*). Less well known is that during neurodevelopment, astrocytes can express positional guidance cues, such as semaphorin 3a, that are required for neuronal circuit formation, through mediating the attraction or repulsion of the growth cone at the axonal tip(*101*). In our gene-based association analysis, *SEMA3A* was the most significantly associated individual gene with brain-wide fiber tract connectivity in the whole genome. Taken together, our data suggest that the formation of fiber tracts in the developing human brain may be affected substantially by positional cues provided by astrocytes, in addition to neurons.

As regards microglia – these phagocytic cells are well known for immune functions but also help to remove dying neurons and prune synapses, as well as modulate neuronal activity(*102*). Less is known of their roles during development, but embryonic microglia are unevenly distributed in the brain and associate with developing axons, which again suggests roles in regulating axonal growth and positional guidance(*103*). Mouse brains without microglia, or with immune activated microglia, show abnormal dopaminergic axon outgrowth(*104*), while disruption of microglial function or depletion of microglia results in a failure of growing axons to adhere and form bundles in the corpus callosum – the largest fiber tract of the brain(*105*). Our data support such observations, through showing that genes up-regulated in microglia in the embryonic human brain are enriched for variants that associate with individual differences in adult white matter connectivity. Further research on the roles of astrocytes and microglia in fiber tract development is therefore warranted.

While our results point especially to genes involved in neurodevelopment, it is also likely that some genetic effects on white matter connectivity act later in life. For example, astrocytes and microglia may affect the maintenance and aging of brain fiber tracts during adulthood, with implications for brain disorders, and possibly suggesting therapeutic targets. We mapped the multivariate associations of polygenic scores for various brain-related disorders and behavioural traits with regional white matter connectivities, and annotated the resulting brain maps using meta-analyzed functional imaging data. Some maps and their annotations were consistent with expectations – for example, polygenic disposition to bipolar disorder was associated with white matter connectivity of brain regions prominently involved in mood, while polygenic dispositions to attention deficit/hyperactivity disorder or autism were associated with the connectivity of regions important for executive functions. Polygenic scores for left-handedness and for schizophrenia were associated with the connectivity of language-related regions – consistent with altered left-hemisphere functional dominance for language in both of these traits(*25, 106*), and a phenotypic association between them(*107*). Polygenic scores for left-handedness and schizophrenia have also been associated with altered structural asymmetry of grey matter in language-related regions(*21, 78*).

Regarding genetic risks for neurological disorders, polygenic scores for Alzheimer’s disease and amyotrophic lateral sclerosis were associated with the connectivity of regions important for working memory, while polygenic scores for epilepsy were associated with connectivity of the default mode network – a set of brain regions involved in internally-initiated thoughts, and semantic and episodic memory(*108*). Previous analysis of white matter tracts in Alzheimer’s disease has indicated a broad-based reduction of connectivity(*109*), so it was striking that the majority of brain regions in the UK Biobank adult population dataset showed increased connectivity with higher polygenic risk for this disorder, even while some core regions of pathology showed decreased connectivity as expected. A similarly notable pattern was seen for amyotrophic lateral sclerosis, where increased polygenic risk was associated with increased structural connectivity for a majority of brain regions. It may be that increased connectivity of some regions occurs as a compensatory re-configuration in response to decreased connectivity of others(*110*). In addition, the UK Biobank volunteer sample is healthier than the general population(*111*). Those at higher polygenic risk who manifest a given disorder may tend not to participate, leaving an unusually healthy set of volunteers among those with high polygenic risk. Such recruitment bias may influence brain correlates of polygenic risk in the UK Biobank.

The brain-wide multivariate GWAS approach that we used provided high statistical power to detect relevant genomic loci, compared to a mass univariate approach(*30*). At the same time, the multivariate results could be queried *post hoc* to identify loci associated with particular tract-wise connectivities of interest. We illustrated this by querying the results with respect to six tracts linking four core regions of the left-hemisphere language network – together approximating to Broca’s and Wernicke’s classically defined functional areas(*112*). Twenty-six implicated loci included the *EPHA3* locus, encoding an ephrin receptor subunit that acts a positional guidance cue for the formation of axon projection maps, and has also been associated with functional connectivity between regional components of the language network that are especially involved in semantics(*69*). This is therefore a concordant genetic finding with respect to both structural and functional connectivity of the human brain’s language network.

We found the heritability of white matter left-right asymmetry metrics to be generally lower than the corresponding unilateral measures from which they were derived, and accordingly many fewer genetic loci were identified in our multivariate GWAS analyses of asymmetry metrics at both node and edge levels. This overall pattern has been found before with respect to other aspects of brain structure and their asymmetries(*20, 35*). One general explanation may be that asymmetry indexes are affected by measurement error in both of the unilateral metrics that are used to calculate them – this may then contribute to larger proportional estimates of non-genetic variance in heritability analysis. It is also likely that asymmetries are affected by a relatively high degree of random developmental variation – the noise inherent in creating complex organs from genomic components(*20, 113, 114*). Nonetheless, we found some of the regional connectivity asymmetries and tract-wise connectivity asymmetries to be significantly heritable. This is consistent with the existence of genetically-regulated, lateralized developmental biases that ultimately give rise to hemispheric specializations, such as left-hemisphere language dominance. Specific loci that we found associated with white matter asymmetries implicated genes involved in synaptic adhesion, glioblastomas, and cytoskeleton modeling.

This study had some limitations: i) We maximized our statistical power for GWAS by using the available data as one large discovery sample, but this did not permit a discovery-replication design(*115*). Nonetheless, ultimately the total combined analysis in the largest available sample is the most representative of the available evidence for association. As mentioned earlier in this section, the various enrichment analyses indicated biological validity of the GWAS findings. Indeed, it has been argued that discovery-replication designs have less utility in the current era of Biobank-scale genetic studies than they used to, and that other forms of validation such as biological enrichment should be given increased weight in interpretation(*116*). ii) We used deterministic tractography which we found to be computationally feasible in more than 30,000 individuals (and which took several months of processing on a cluster server). An alternative approach - probabilistic tractography - may have advantages insofar as it permits modelling of multiple tract orientations per voxel(*117, 118*), although run times are generally higher, and our approach yielded genetic results with clear biological validity for white matter tracts. iii) This was a large-scale observational mapping study, which meant that many of the analyses were screen-based and descriptive. Science proceeds through a combination of observation and hypothesis testing – this study incorporated both to varying degrees. Some of the biological observations were striking and informative, for example the likely involvements of microglia and astrocytes in affecting white matter tracts during embryonic and fetal development, which should now be studied more extensively in animal models. iv) This study did not consider rare genetic variants (with population frequencies below 1%). Future analysis of the UK Biobank’s exome and genome sequence data in relation to white matter connectivity may reveal further genes and suggest additional mechanisms, cell types and lifespan stages in affecting inter-individual variation.

In summary, we used large-scale analysis to chart the white matter connectivity of the human brain, its multivariate genetic architecture, and its associations with polygenic dispositions to brain-related disorders and behavioural traits. The analyses implicated specific genomic loci, genes, pathways, cell types, developmental stages, brain regions, fiber tracts, and cognitive functions, thus integrating multiple levels of analysis, and suggesting a range of future research directions at each of these levels.

## Materials and Methods

### Sample quality control

This study was conducted under UK Biobank application 16066, with Clyde Francks as principal investigator. The UK Biobank received ethical approval from the National Research Ethics Service Committee North West-Haydock (reference 11/NW/0382), and all of their procedures were performed in accordance with the World Medical Association guidelines(*119*). Written informed consent was provided by all of the enrolled participants. We used the dMRI data released in February 2020, together with the genome-wide genotyping array data. For individuals with available dMRI and genetic data, we first excluded subjects with a mismatch of their selfreported and genetically inferred sex, with putative sex chromosome aneuploidies, or who were outliers according to heterozygosity (principle component corrected heterozygosity >0.19) and genotype missingness (missing rate >0.05) as computed by Bycroft et al(*120*). To ensure a high degree of genetic homogeneity, analysis was limited to participants with white British ancestry, which was defined by Bycroft et al.(*120*) using a combination of self-report and cluster analysis based on the first six principal components that capture genetic ancestry. We also randomly excluded one subject from each pair with a kinship coefficient >0.0442, as calculated by Bycroft et al.(*120*). This inclusion procedure finally resulted in 30,810 participants, with a mean age of 63.84 years (range 45-81 years), 14,636 were male and 16,174 were female.

### Genetic quality control

We downloaded imputed SNP and insertion/deletion genotype data from the UK Biobank (i.e. v3 imputed data released in March 2018). QCTOOL (v.2.0.6) and PLINK v2.0(*121*) were used to perform genotype quality control. Specifically, we excluded variants with minor allele frequency <1%, Hardy-Weinberg equilibrium test p value <1×10^−7^ and imputation INFO score <0.7 (a measure of genotype imputation confidence), followed by removing multi-allelic variants that cannot be handled by many programs used in genetic-related analyses. This pipeline finally yielded 9,803,735 bi-allelic variants.

### Diffusion MRI-Based Tractography

Diffusion MRI data were acquired from Siemens Skyra 3T scanners running protocol VD13A SP4, with a standard Siemens 32-channel RF receive head coil(*122*). We downloaded the quality-controlled dMRI data which were preprocessed by the UK Biobank brain imaging team(*122, 123*) (UK Biobank data field 20250, first imaging visit). The preprocessing pipeline included corrections for eddy currents, head motion, outlier slices, and gradient distortion. We did not make use of imaging-derived phenotypes released by the UK Biobank team, such as FA and mean diffusivity. Rather, we used the quality-controlled dMRI data to perform tractography in each individual, which generated three-dimensional curves that characterize white matter fiber tracts. Briefly, diffusion tensors were modeled to generate a FA image in native diffusion space, which was used for deterministic diffusion tensor tractography using MRtrix3(*124*). Streamlines were seeded on a 0.5 mm grid for every voxel with FA 0.15 and propagated in 0.5 mm steps using fourth-order Runge-Kutta integration. Tractography was terminated if the streamline length was <20 or >250 mm, if it turned an angle >45°, or reached a voxel with an FA <0.15. These parameters were consistent with a previous study exploring the structural network correlates of cognitive performance using the UK Biobank dataset(*125*). Tens of thousands of streamlines were generated to reconstruct the white matter connectivity matrix of each individual on the basis of the Automated Anatomical Labelling atlas(*26*) comprising a total of 90 regions encompassing cortical and subcortical structures (45 regions per hemisphere). This deterministic tractography process took roughly 16 weeks on 6 cluster server nodes running in parallel.

We calculated the Euclidean distance between the centroids of each pair of connected regions, according to brain standard space (MNI template provided with the Automated Anatomical Labeling atlas(*26*)), to index the relative physical distances between regions.

### Network construction and analysis

Describing the structural network of each participant requires the definition of network nodes and edges. In this study, the network nodes corresponded to the 90 regions of the Automated Anatomical Labeling atlas(*26*). The labeling system integrates detailed anatomical features from sulcal and gyral geometry, reducing anatomical variability that can arise from spatial registration and normalization of brain images taken from different individuals(*26*). For each participant, the T1 images were nonlinearly transformed into the ICBM152 T1 template in the MNI space to generate the transformation matrix(*126*). Inverse transformation was used to warp the Automated Anatomical Labeling atlas(*26*) from the MNI space to native space. Discrete labeling values were preserved using a nearest-neighbor interpolation method(*126*). Two nodes were considered connected if they were joined by the endpoints of at least one reconstructed streamline. Network edges were computed by the number of streamlines connecting a given pair of regions, while dividing by the volume of the two regions, because regions with larger volumes tend to have more streamlines connecting to them. We only included edges that were detected in at least 80% of participants. This yielded a zero-diagonal symmetrical 90×90 undirected connectivity matrix for each participant, in which 947 edges were retained. The regional connectivity of a node was then defined as the sum of all existing edges between that node and all other nodes in the network, reflecting the importance of that node in the overall network.

Asymmetry of network measures at the node and edge levels was assessed by the asymmetry index (AI) for each pair of network measures with left and right hemisphere homologues, using the following formula per individual:

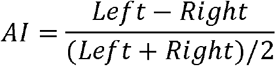

In this formula, the denominator normalizes by the bilateral measure. Positive values of the AI represent leftward asymmetry (greater left than right), and vice versa. If the corresponding left and right measures for a given individual and homologous pair were both zero, we set the AI as zero.

Rank-based inverse normalization across individuals was performed on each network measure, and regression on age, nonlinear age (i.e. (age-mean_age)^2^), assessment center, genotype measurement batch and sex. Residuals were then further regressed on the first ten genetic principal components that capture population genetic diversity(*120*), followed by rank-based inverse normalization of the residuals once more. These normalized, transformed measures were used for subsequent genetic analyses.

### SNP-based heritability

We constructed a genetic relationship matrix using 9,516,306 variants on the autosomes with minor allele frequencies >1%, INFO score >0.7 and Hardy-Weinberg equilibrium p value >1×10^−7^, using GCTA(*29*) (version 1.93.0beta). We further excluded one random participant from each pair having a kinship coefficient higher than 0.025 (as SNP-based heritability analysis is especially sensitive to participants with higher levels of relatedness), yielding 29,027 participants for this particular analysis. Genome-based restricted maximum likelihood analyses were then performed to estimate the SNP-based heritability for each normalized structural network measure, again using GCTA(*29*). Bonferroni correction was applied separately for each type of network measure to identify those that were significantly heritable at adjusted p<0.05: 90 regional connectivities, 851 tract-wise connectivities, 15 regional connectivity AIs and 107 tract-wise connectivity AIs were significantly heritable.

### Multivariate genome-wide association analysis

A total of 9,803,735 bi-allelic variants were used for mvGWAS analysis, spanning all autosomes and chromosome X. The sample size for mvGWAS was 30,810 (see sample quality control, above). We applied the multivariate omnibus statistical test (MOSTest) toolbox(*30*) to perform mvGWAS analysis for the significantly heritable measures, separately for regional connectivities, tract-wise connectivities, regional connectivity AIs, and tract-wise connectivity AIs (therefore four mvGWAS analyses in total). MOSTest can leverage the distributed nature of genetic influences across hundreds of spatially distributed brain phenotypes, while accounting for their covariances, which can boost statistical power to detect variant-phenotype associations(*30*). Specifically, the multivariate correlation structure is determined on randomly permuted genotype data. MOSTest calculates the Mahalanobis norm as the sum of squared de-correlated z-values across univariate GWAS summary statistics, to integrate effects across measures into a multivariate z statistic for each genetic variant, and uses the gamma cumulative density function to fit an analytic form for the null distribution. This permits extrapolation of the null distribution below the p=5×10^−8^ significance threshold without performing an unfeasible number of permutations (5 ×10^−8^ is a widely used threshold for GWAS multiple test correction in European-descent populations(*127, 128*)). Close matching of the null p value distributions from the permuted and analytic forms indicate that the method correctly controls type 1 error – this was the case for all four of our mvGWAS analyses (Supplementary Figures 4, 5, 12 and 13). In this framework the signs (positive or negative) of univariate z scores indicate the corresponding directions of effects (with respect to increasing numbers of minor alleles at a given SNP), whereas multivariate z scores are always positive.

### Identification of genomic loci and functional annotations

We used FUMA (version v1.3.7)(*31*) to identify distinct genomic loci showing significant multivariate associations with brain structural connectivity, and apply functional annotations, using default parameters. Linkage disequilibrium (LD) structure was applied according to the 1000 Genomes European reference panel(*129*). SNPs with genome-wide significant mvGWAS p values <5×10^−8^ that had LD r^2^<0.6 with any others were identified. For each of these SNPs, other SNPs that had r^2^−0.6 with them were included for further annotation (see below), and independent ‘lead SNPs’ were also defined among them as having low LD (r^2^<0.1) with any others. If LD blocks of significant SNPs were located within 250□kb of each other, they were merged into one genomic locus. Therefore, some genomic loci could include one or more independent lead SNPs. The major histocompatibility complex region on chromosome 6 was excluded from this process by default, because of its especially complex and long-range LD structure.

Functional annotations were applied by matching chromosome location, base-pair position, reference and alternate alleles to databases containing known functional annotations, which were ANNOVAR(*130*) categories, Combined Annotation-Dependent Depletion(*131*) scores, RegulomeDB(*132*) scores and chromatin state(*133, 134*):

1. ANNOVAR catogorizes SNPs on the basis of their locations with respect to genes, e.g. exonic, intronic and intergenic, using Ensembl gene definitions.
2. Combined Annotation-Dependent Depletion scores predict deleteriousness, with scores higher than 12.37 suggesting potential pathogenicity(*135*).
3. RegulomeDB scores integrate regulatory information from eQTL and chromatin marks, and range from 1a to 7, with lower scores representing more importance for regulatory function.
4. Chromatin states show the accessibility of genomic regions, and were labelled by 15 categorical states on the basis of five chromatin marks for 127 epigenomes in the Roadmap Epigenomics Project(*134*), which were H3K4me3, H3K4me1, H3K36me3, H3K27me3 and H3K9me3. For each SNP, FUMA calculated the minimum chromatin state across 127 tissue/cell-types in the Roadmap Epigenomics Project(*120*). Categories 1-7 are considered open chromatin states.

We also used FUMA to annotate significantly associated SNPs, and other candidate SNPs that had r^2^≥0.6 with them, according to previously reported phenotype associations (p<5□×10^−5^) in the National Human Genome Research Institute-European Bioinformatics Institute catalogue(*32*).

### Multivariate association profiles of independently associated lead SNPs

For each SNP, MOSTest derives a z-score for each brain measure, calculated from the p value of the univariate association of that SNP with each individual measure. The z-scores give an indication of which measures contribute most to the multivariate association for a given SNP(*30*). We used the z-scores from the mvGWAS of fiber tracts to identify lead SNPs that were significantly associated with at least one from a set of six left-hemisphere language-related fiber tracts (see main text). To determine significance in this context, a threshold z-score with unsigned magnitude >3.7 was applied, corresponding to a p value of 2.16×10^−4^ (i.e. p<0.05 after Bonferroni correction for all 231 lead SNPs from the mvGWAS of fiber tracts, and considering six fiber tracts).

To determine which structural connectivity measures contributed most to the multivariate associations as considered across lead SNPs, we summed the unsigned univariate z-scores separately for each measure across all lead SNPs (separately for the mvGWAS analyses of regional connectivities and fiber tracts).

### SNP-to-gene mapping

Independent lead SNPs, and candidate SNPs having LD r^2^>0.6 with a lead SNP, were mapped to genes in FUMA using the following three strategies.

1. Positional mapping was used to map SNPs to protein-coding genes based on physical distance (within 10□kb) in the human reference assembly (GRCh37/hg19).
2. eQTL mapping was used to annotate SNPs to genes up to 1□Mb away based on a significant eQTL association, i.e. where the expression of a gene is associated with the allelic variation, according to information from four brain-expression data repositories, including PsychENCORE(*52*), CommonMind Consortium(*48*), BRAINEAC(*136*) and GTEx v8 Brain(*49*). FUMA applied a FDR of 0.05 within each analysis to define significant eQTL associations.
3. Chromatin interaction mapping was used to map SNPs to genes on the basis of seven brain-related Hi-C chromatin conformation capture datasets: PsychENCORE EP link (one way)(*52*), PsychENCORE promoter anchored loops(*48*), HiC adult cortex(*55*), HiC fetal cortex(*55*), HiC (GSE87112) dorsolateral prefrontal cortex(*50*), HiC (GSE87112) hippocampus(*50*) and HiC (GSE87112) neural progenitor cells(*50*). We further selected only those genes for which one or both regions involved in the chromatin interaction overlapped with a predicted enhancer or promoter region (250□bp upstream and 500□bp downstream of the transcription start site) in any of the brain-related repositories from the Roadmap Epigenomics Project(*134*), that is; E053 (neurospheres) cortex, E054 (neurospheres) ganglion eminence, E067 (brain) angular gyrus, E068 (brain) anterior caudate, E069 (brain) cingulate gyrus, E070 (brain) germinal matrix, E071 (brain) hippocampus middle, E072 (brain) inferior temporal lobe, E073 (brain) dorsolateral prefrontal cortex, E074 (brain) substantia nigra, E081 (brain) fetal brain male, E082 (brain) fetal brain female, E003 embryonic stem (ES) H1 cells, E008 ES H9 cells, E007 (ES-derived) H1 derived neuronal progenitor cultured cells, E009 (ES-derived) H9 derived neuronal progenitor cultured cells and E010 (ES-derived) H9 derived neuron cultured cells. FUMA applied a FDR of 1 × 10^−6^ to identify significant chromatin interactions (default parameter), separately for each analysis.

### Gene-based association analysis

MAGMA (v1.08)(*37*), with default parameters as implemented in FUMA (SNP-wise mean model), was used to test the joint association arising from all SNPs within a given gene (including 50□kb upstream to 50□kb downstream), while accounting for LD between SNPs. SNPs were mapped to 20,146 protein-coding genes on the basis of National Center for Biotechnology Information build 37.3 gene definitions, and each gene was represented by at least one SNP. Bonferroni correction was applied for the number of genes (p<0.05/20,146), separately for each mvGWAS.

### Gene-set enrichment analysis

MAGMA (v1.08), with default settings as implemented in FUMA, was used to examine the enrichment of association for predefined gene sets. This process tests whether gene-based p values among all 20,146 genes are lower for those genes within pre-defined functional sets than the rest of the genes in the genome, while correcting for other gene properties such as the number of SNPs. A total of 15,488 gene sets from the MSigDB database version 7.0(*61*) (5500 curated gene sets, 7343 gene ontology (GO) biological processes, 1644 GO molecular functions, and 1001 GO cellular components) were tested. Bonferroni correction was applied to correct for the number of gene sets (p<0.05/15,488), separately for each mvGWAS.

### Cell-type-specific expression analysis in developing human cortex

Based on a linear regression model, the CELL TYPE function of FUMA was used to test whether gene-based association z-scores were positively associated with higher expression levels in certain cell types, based on singlecell RNA sequencing data from the developing human prefrontal cortex (GSE104276)(*63*). This dataset comprised 1) expression per cell type per age group, ranging from 8 to 26 postconceptional weeks, and 2) expression profiles per cell type, averaged over all ages combined. Results were considered significant if the association p values were smaller than the relevant Bonferroni threshold for the number of cell types/age groups. Analysis was performed separately for each mvGWAS.

### Developmental stage analysis

We used MAGMA (default settings as implemented in FUMA) to examine whether lower gene-based association p values tended to be found for genes showing relatively higher expression in BrainSpan gene expression data(*62*) from any particular lifespan stages when contrasted with all other stages, separately for 29 different age groups ranging from 8 postconceptional weeks to 40 years old, and 11 defined developmental stages from early prenatal to middle adulthood. A FDR threshold of 0.05 was applied separately for each analysis. Positive beta coefficients for this test indicate that genes showing more evidence for association are relatively upregulated on average at a given lifespan stage.

### Polygenic disposition to brain-related disorders or behavioural traits

We used PRS-CS(*76*) to compute polygenic scores for 30,810 UK Biobank individuals (see Sample quality control) for each of the following brain-related disorders or behavioural traits, using GWAS summary statistics from previously published, large-scale studies: schizophrenia (n=82,315)(*77*), bipolar disorder (n=51,710)(*80*), autism (n=46,350)(*83*), attention deficit/hyperactivity disorder (n=55,374)(*84*),, left-handedness (n=306,377)(*21*), Alzheimer’s disease (n=63,926)(*87*), amyotrophic lateral sclerosis (n=152,268)(*90*) and epilepsy (n=44,889)(*93*). None of these previous studies used UK Biobank data, except for the GWAS of left-handedness(*21*) – however the individuals in that GWAS were selected to be non-overlapping and unrelated to those with brain image data from the February 2020 data release, so that none of the 30,810 UK Biobank individuals from the present study had been included in that GWAS. This ensured that training and target sets for polygenic score calculation were independent. PRS-CS infers posterior effect sizes of autosomal SNPs on the basis of genome-wide association summary statistics, within a high-dimensional Bayesian regression framework. We used default parameters and the recommended global effect size shrinkage parameter φ=0.01, together with linkage disequilibrium information based on the 1000 Genomes Project phase 3 European-descent reference panel(*137*). Polygenic scores were calculated using 1,097,390 SNPs for schizophrenia, 1,098,372 SNPs for bipolar disorder, 1,092,080 SNPs for autism, 1,042,054 SNPs for attention deficit/hyperactivity disorder, 1,103,632 SNPs for left-handedness, 1,105,067 SNPs for Alzheimer’s disease, 1,085,071 SNPs for amyotrophic lateral sclerosis, and 852,343 SNPs for epilepsy (these numbers came from 3-way overlaps between UK Biobank data, GWAS results, and 1000 Genomes data). PRS-CS has been shown to perform in a highly similar manner to other established polygenic risk algorithms, with noticeably better out-of-sample prediction than an approach based on p value thresholds and LD clumping(*138, 139*).

Polygenic scores were linearly adjusted for the effects of age, nonlinear age (i.e. (age-mean_age)^2^), assessment centre, genotype measurement batch, sex, and the first ten genetic principal components that capture population genetic diversity, before performing rank-based inverse normalization (i.e. the same set of covariate effects that the brain metrics were adjusted for - see Network construction and analysis). These adjusted and normalized polygenic scores were used as input for subsequent analyses.

Separately for polygenic scores for each disorder or behavioural trait, canonical correlation analysis across 30,810 participants (‘*canoncorr*’ function in MATLAB) was used to test multivariate association with the 90 heritable regional connectivity measures (which had also been adjusted for covariates and normalized - see Network construction and analysis). This multivariate analysis identified a linear combination of the 90 regional connectivity measures (i.e. a canonical variable) that maximally correlated with the polygenic score for a particular disorder or behavioural trait across participants. Separately for the polygenic score of each disorder or behavioural trait, the cross-participant Pearson correlation between each regional connectivity and the canonical variable was used as a loading, reflecting the extent and direction of the contribution that a regional connectivity made to a particular multivariate association.

We also assessed the pairwise correlations across individuals between adjusted and normalized polygenic scores for the different disorders and behavioural traits.

As the *APOE* locus is known to have a substantial effect on the risk of Alzheimer’s disease, we also re-calculated polygenic scores for this disease after excluding a region from Chr19:45,116,911 to Chr19:46,318,605 (GRCh37)(*140*) around this locus, and repeated the residualization, normalization and canonical correlation analyses to check that the results stably reflected the polygenic contribution to risk.

### Functional annotation of brain regions associated most strongly with polygenic scores

From each separate canonical correlation analysis of polygenic scores and regional connectivity, we identified the regions showing loadings of >0.2 or <-0.2, which were then used to define a single mask in standard brain space (Montreal Neurological Institute space 152) (i.e. one mask for each polygenic score). Each mask was analyzed using the ‘decoder’ function of the Neurosynth database (http://neurosynth.org), a platform for large-scale synthesis of functional MRI data(*99*). This database defines brain-wide activation maps corresponding to specific cognitive or behavioural task terms using meta-analyzed functional activation maps. The database included 1,334 term-specific activation maps corresponding to cognitive or behavioural terms from 14,371 studies. Each mask that we created was used separately as input to define a brain-wide co-activation map based on all studies in the database. The resulting co-activation maps were then correlated with each of the 1,334 term-specific activation maps(*99*). We report only terms with correlation coefficients r>0.2, while excluding anatomical terms, nonspecific terms (e.g., ‘Tasks’), and one from each pair of virtually duplicated terms (such as ‘Words’ and ‘Word’). This method does not employ inferential statistical testing, but rather ranks terms based on the correlations between their activation maps and that of the input mask.

## Data Availability

The primary data used in this study are available via the UK Biobank website www.ukbiobank.ac.uk. Other publicly available data sources and applications are cited in the Methods section. The GWAS summary statistics are made available online within the GWAS catalog https://www.ebi.ac.uk/gwas/.

## Code availability

This study used openly available software and codes, specifically GCTA (https://cnsgenomics.com/software/gcta/#GREML), MRtrix3 (https://www.mrtrix.org/), MOSTest (https://github.com/precimed/mostest), FUMA (https://fuma.ctglab.nl/), MAGMA (https://ctg.cncr.nl/software/magma, also implemented in FUMA), PRS-CS (https://github.com/getian107/PRScs) and Neurosynth (https://www.neurosynth.org/).

## Author contributions

ZS: Conceptualization, methodology, analysis, visualization, original draft writing, review, and editing. DS: Methodology, analysis, review, and editing. SEF: Methodology, review, and editing. CF: Conceptualization, direction, supervision, original draft writing, review, and editing.

## Disclosures

The authors declare no competing interests.

## Acknowledgements

This research was funded by the Max Planck Society (Germany), and conducted using the UK Biobank resource under application no. 16066 with C.F. as the principal applicant. Our study made use of quality-controlled brain images generated by an image-processing pipeline developed and run on behalf of UK Biobank. The funders had no role in study design, data collection and analysis, the decision to publish or preparation of the manuscript.

## Notes

### Competing Interest Statement

The authors have declared no competing interest.

### Summary of Updates

As requested by the journal, the original version of the paper should be uploaded to bioRxiv.

